# Fragmentation of genomic DNA using Covaris g-Tubes prior to library preparation for sequencing

**DOI:** 10.1101/2024.05.29.596380

**Authors:** Nick Crang, Orlando Contreras-López

**Affiliations:** National Genomics Infrastructure (NGI) – SciLifeLab, Science for Life Laboratory, Tomtebodavägen 23b, 171 65 Solna, Sweden; Skolan för Kemi, Bioteknologi och Hälsa, Teknikringen 30, 114 28 Stockholm, Sweden

**Keywords:** DNA, Oxford Nanopore Sequencing, Long Read Sequencing, Library Preparation

## Abstract

Covaris g-TUBEs can be used to fragment DNA to pre-determined sizes based on the relative centrifugal force that they are run at. They are recommended for use while preparing Oxford Nanopore Technology libraries by the manufacturer. However, the volumes and DNA concentration typically used for ONT libraries are outside the range of the example data provided by Covaris. Here, we ran g-TUBEs at three different relative centrifugal forces and determined the effect on DNA fragmentation in the range 0.5 - 4 µg. This dataset can be used to inform the effective fragmentation of DNA for creating Oxford Nanopore libraries of an optimal size.

## Introduction

It can sometimes be beneficial to fragment genomic DNA when preparing libraries for sequencing on the Oxford Nanopore Technologies (ONT) sequencers. Fragmenting the DNA increases the molarity of the sample, along with making the length distribution more uniform and allowing for a more accurate estimation of sample molarity. The decrease in fragment length also increases the yield of the ONT library preparation as longer fragments are more easily lost during clean-up steps. This is compounded by the increased ease of sequencing of shorter fragments, which will result in a greater sequencing yield for the library. However, fragmentation does decrease the N50 of the library (defined as the largest length L such that 50% of all nucleotides are contained in contigs of size at least L^1^) and the length of the sequence reads is one of the great virtues of ONT sequencing.

There are a number of ways to fragment DNA prior to library preparation, but one of the quickest is through use of Covaris g-TUBEs. Here, DNA is fragmented by passing the sample through a fixed aperture membrane using bench top centrifugation resulting in DNA fragmentation with predictable fragment sizing. The relative centrifugal force (RCF) used to pass the sample through the aperture can be varied resulting in greater or lesser mechanical shearing force upon the DNA and thus fragmenting it into shorter or longer fragments.

According to the manufacturers of the Covaris g-TUBEs, the three factors that can affect the fragmentation are:

1. Sample volume
2. DNA concentration
3. RCF applied to the sample

It is worth noting that when fragmenting DNA to sizes <8 kbp, the processing time is reduced to 30 seconds from 1 minute (*Table 1*). This is not discussed in the literature available from the manufacturer, and since this fragment size is much shorter than the 20 kbp ideal we aimed to achieve, we decided to maintain a 1 minute processing time for all samples.

**Table 1.**
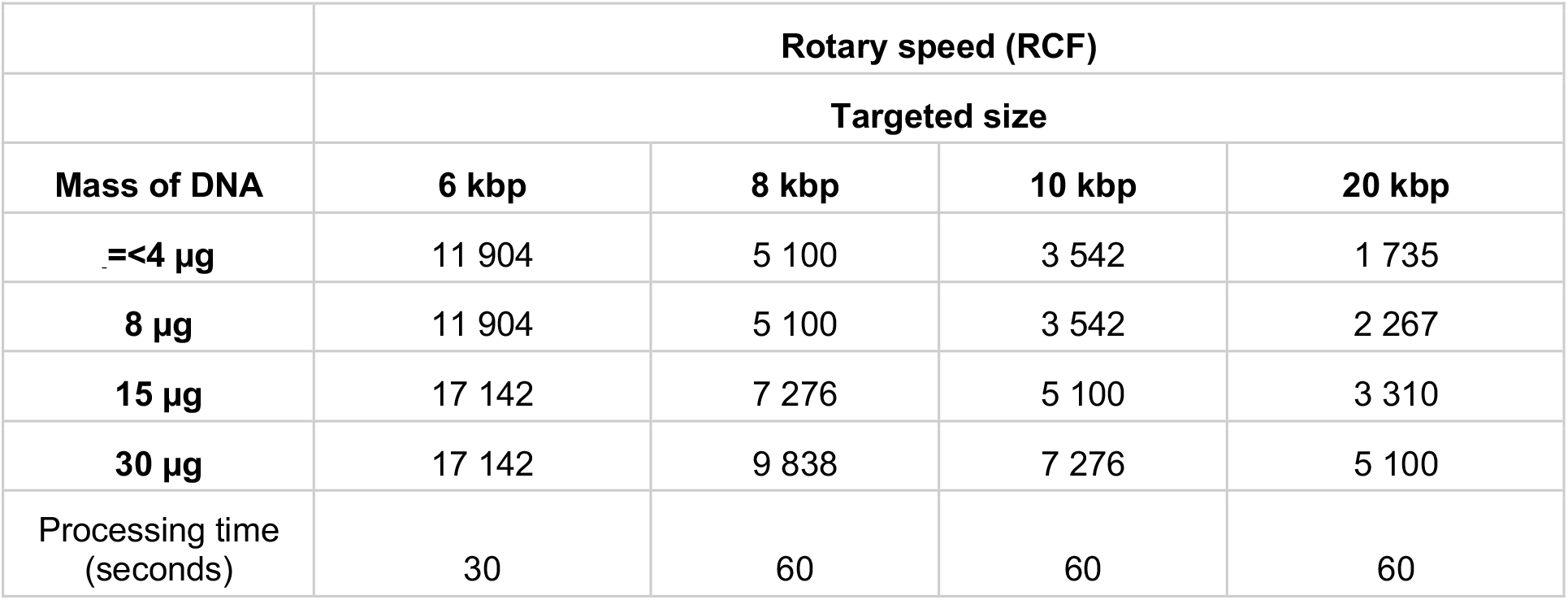
Modified after the g-TUBE user manual (pn_010154 rev D) from Covaris A table. The RCF required to fragment DNA to different lengths for different DNA concentrations is indicated. These values are for a sample volume of 150 µL. NOTE the original table was provided in RPM, this has been converted to RCF for ease of comparison using the formula RCF = 1.118 * 10^(-5) * r * RPM^2

We typically have a concentration of DNA around 1-2 µg provided by users for preparation of libraries for ONT. The volume we used for fragmentation is typically 50 µL rather than the 150 µL suggested by Covaris as this means we do not have to include a DNA concentration step to reduce our fragmented DNA to a usable sample volume for library input. We have previously observed this smaller volume and DNA concentration has a measurable effect on fragmentation, with an RCF of 1 500 resulting in an average fragment size of between 14 051 bp when using 50 µL of 0.5 µg sample and 15 651 bp when using 50 µL of 1 µg sample. Both values are considerably smaller than the 20 kbp suggested by *Table 1*. As the table provided by Covaris gave unified recommendations for samples containing <4 µg DNA, we performed a fragmentation test using multiple concentrations within our expected sample range (0.5 and 4 µg).

In this study, we have two primary aims:

1. Identify the RCF(s) most appropriate to achieve 20 kb fragmentation of DNA within our operating range (0.5–4 µg in 50 µl)
2. Identify the most appropriate method for size determination of large fragments (DNF-464 HS Large Fragment Kit for Fragment Analyzer or FP-1002 Genomic DNA 165 kb for Femto Pulse)

We have additionally determined that an individual Covaris g-TUBE, though described as single use, can in fact be reused multiple times, though our observation is that membrane clogging typically becomes a problem after four sample preps.

It was decided to test three replicates each for the DNA amounts 0.5, 1, 2 and 4 µg at 1000, 1500 and 2000 RCF. All fragmentation tests were performed for 1 minute in 50 µL. The degree of fragmentation was measured in both the Femto Pulse System (Agilent), hereafter termed FEMTO, and the 5200 Fragment Analyzer System (Agilent), hereafter termed FA instruments.

The kit used with the FEMTO should have greater resolving power for fragments above 20 kb. The FEMTO has both a short and extended run settings which can be used to enhance the sample resolution, though this increases the run time from 40 minutes to 3 hours per sample.

## Material and methods

### Dilution

Commercially sourced human DNA (#G1471; Promega Corporation) was diluted using nuclease-free water to prepare four sample dilution pools containing 0.5, 1, 2 and 4 µg of DNA / 50 µL of volume. Each of these pools were 500 µL in total volume, and were used to provide replicate samples for each test condition. Pipette mixing of the pools was performed using wide bore pipette tips to prevent the high molecular weight DNA being damaged.

### DNA fragmentation

A total volume of 50 µL was loaded into a single g-TUBE #520079 (Covaris; Perkin Elmer). Each g-Tube was used for a single RCF and reused four times, with samples being loaded in the order 0.5, 1, 2 and finally 4 µg of DNA. Fragmentation was achieved via one minute centrifugation at the specified RCF. Samples were run on a Centrifuge 5430 (Eppendorf).

### Fragment Analyzer

Samples were diluted to within the working concentration of the assay with nuclease free water. Two µL of each diluted sample were run using the HS Large Fragment 50kb Kit #DNF-464-0500 (Agilent) according to the manufacturer’s instructions on the 5200 Fragment Analyzer System # M5310AA, 12 Capillaries, 33 cm. Data was analysed using PROSize v5.0 software (Agilent).

### Femto Pulse

Samples were diluted to within the working concentration of the assay with nuclease-free water. Samples were run in both short and extended run formats on the Femto Pulse System, F12218, 12 Capillaries, 33 cm #SN F12218). Two µL of each diluted sample were run using the gDNA 165kb Analysis Kit #FP-1002-0275 (Agilent) according to the manufacturer’s instructions. Data was analysed using PROSize v5.0 software (Agilent).

## Results

The sample preparation via the g-Tubes was conducted without incident, and fragmentation was clearly visible on the traces from both the FA and FEMTO (*Figure 1, 3 Supplementary Figure 1*).

**Fig 1.**
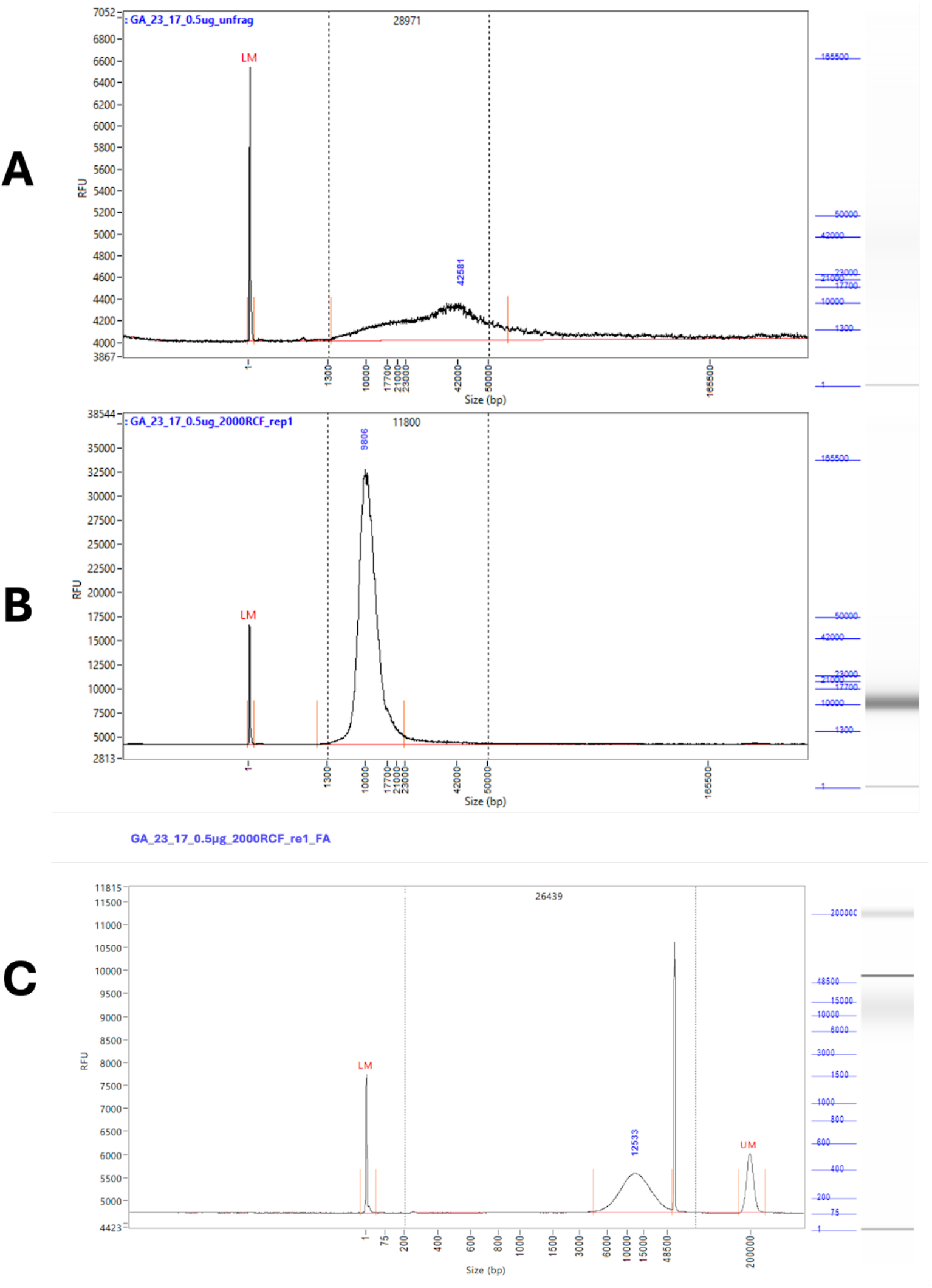
Representative traces from extended run Femto Pulse. A) Shows unfragmented DNA, diluted to 0.5 µg / 50 µL. The tail of the peak extends well above 50 kb and down to 1.3 kb. The peak height is 42 kb. B) Shows the DNA taken from the original 0.5 µg / 50 µL pool after fragmentation at 2000 RCF. A single clean peak is now visible with a peak height of 9 kb and a tail reduced to significantly less than 50 kb. C) Shows the DNA taken from the original 0.5 µg / 50 µL pool after fragmentation at 2000 RCF run on the FA. The high molecular weight artefact gives a much greater average fragment size despite a peak height of 12.5 kbp

Prior to fragmentation, the trace was characterised by a broad peak with a very long tail up to the resolution limit of 165 000 bp (Figure 1A, 3A). By contrast, after fragmentation a single clean peak was visible with a limited tail ending well below 50 000 bp (Figure 1B). This pattern was consistent across all centrifugation speeds and quantities of DNA fragmented (*Figures 1, 2, 3*).

A problem was encountered when running samples on the FA, with numerous artefacts present during analysis, making it difficult to accurately size the fragments (Supplementary Figure 1). These artefacts appeared in multiple runs at different sizes and lanes, suggesting that they are poorly resolved DNA fragments rather than sample contamination. These artefacts were also seen when using the short run settings on the FEMTO, but were successfully resolved during an extended run. While there are only minor differences overall in the observed fragment sizes between the two instruments, the lack of artefacts from the extended run on the FEMTO gives us greater confidence in the accuracy of the results. The variation between replicates demonstrates the limitations of the g-Tube in terms of accurate fragmentation, with a size range of +/-4 Kb seen between replicates regardless of RCF and DNA quantity. Overall, when sizes could be resolved they were very similar between the FA and FEMTO with an R^2^ value calculated at ≥ 0.8 (*supplementary data*).

We observed a general trend held true, that as the amount of DNA in the sample increased, the average size of the fragments increased for a given RCF (*Figure 2*). This effect was most apparent at 1000 and 1500 RCF, though it decreased as the RCF increased, with little difference between the four DNA concentrations at 2000 RCF.

**Fig 2.**
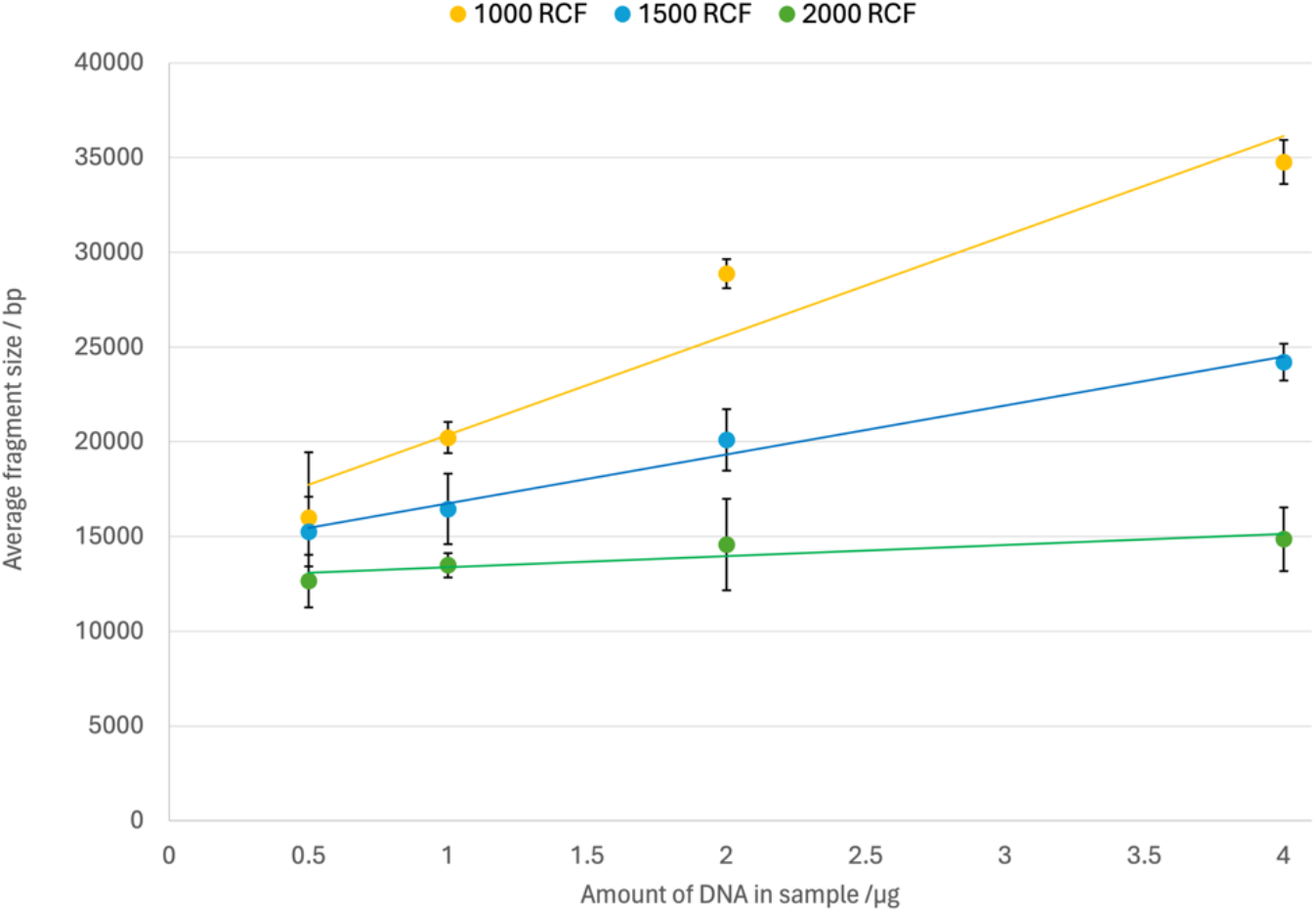
Fragmentation results measured with an extended run on the Femto Pulse # M5330AA (Agilent). Values are an average of three replicates with error bars of standard deviation. The three series indicate the centrifugation speed used during the DNA fragmentation measured in RCF.

It was also observed, as expected, that increasing the RCF reduced the average fragment size. This effect was much more pronounced as the DNA concentration increased, with little difference seen between the three RCFs at 0.5 µg of DNA, where all nine samples were in the range 12-21 000 bp (*Figure 2, Table 2*). By contrast at 4 µg the average size of each fragment at the three RCFs were separated by 10 000 bp, spanning the range 15-35 000 bp (*Figure 2, Table 2*)

**Table 2.** Fragmentation results measured with an extended run on the Femto Pulse # M5330AA (Agilent). Highlighted in **bold** text are RCF conditions that gave optimal fragmentation for that DNA concentration No consistent difference was observed for fragmentation at 2000 RCF, or when the amount of DNA was 0.5 µg, so no values are highlighted.

| Results from FEMTO |  |  |  |
| --- | --- | --- | --- |
| Amount of DNA (µg) | RCF |  |  |
|  | 1000 RCF | 1500 RCF | 2000 RCF |
| 0.5 | 17 445 | 16 756 | 11 800 |
| 0.5 | 14 536 | 13 216 | 11 917 |
| 0.5 | 21 400 | 15 807 | 14 258 |
| 1 | **21 022** | 18 461 | 14 182 |
| 1 | **20 266** | 16 112 | 13 370 |
| 1 | **19 388** | 14 792 | 12 901 |
| 2 | 29 752 | **21 566** | 17 358 |
| 2 | 28 411 | **18 364** | 13 455 |
| 2 | 28 430 | **20 365** | 12 931 |
| 4 | 35 252 | 25 225 | 16 784 |
| 4 | 35 606 | 24 082 | 13 642 |
| 4 | 33 434 | 23 312 | 14 179 |

A clear change was apparent in the traces after g-Tube fragmentation. The broad peak characteristic of the unfragmented DNA (*Figure 1A, Figure 3A*) was replaced with a single, far narrower peak (*Figure 3 B-E*). The breadth of the peak as well as the average fragment size increased along with the amount of DNA in the sample.

**Fig 3.**
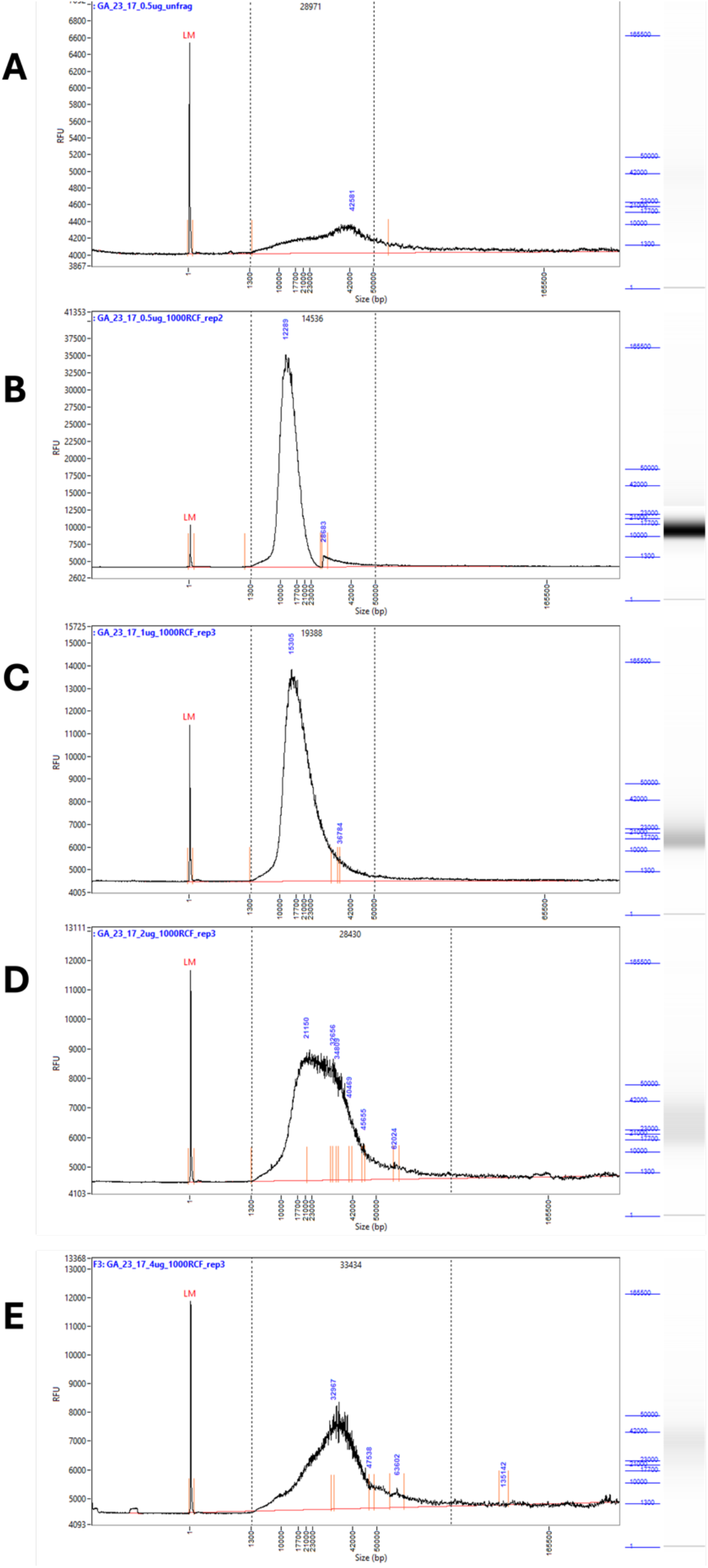
Representative traces of the fragmentation results at RCF 1000 from 0.5 – 4 µg of input DNA. Samples measured with an extended run on the Femto Pulse # M5330AA (Agilent). A) Shows unfragmented DNA, diluted to 0.5 µg / 50 µL. B) Shows DNA diluted to 0.5 µg / 50 µL fragmented at RCF 1000. C) Shows DNA diluted to 1 µg / 50 µL fragmented at RCF 1000. D) Shows DNA diluted to 2 µg / 50 µL fragmented at RCF 1000. E) Shows DNA diluted to 4 µg / 50 µL fragmented at RCF 1000.

## Discussion

On average the correlation between FA and FEMTO results is good (R^2^≥ 0.8, supplementary data), but individual measurements can differ as much as 5000bp between the systems. Based on this observation, if accuracy is needed we would recommend having replicates of the same sample in one run or running the same samples several times. Given the prevalence of unresolvable artefacts during FA and short FEMTO runs we would also recommend performing an extended run on the FEMTO to guarantee high quality usable results. If an approximation of the size is sufficient, then a short FEMTO run or a FA run are equally applicable.

Generally, smaller fragment sizes are given by FEMTO than FA, but this was not always the case. The most pronounced effect was that larger fragments tended to appear even larger when run on FA, which makes sense given the greater resolving power of the kit used with the FEMTO in this range.

It should be noted that sequencing performance of DNA fragmented this way has not been tested during this investigation. Anecdotal reports from other sequencing labs indicates that many more short fragments are left in the samples after using the g-Tube compared to the alternative fragmentation method Meagruptor 3 (Diagenode), suggesting a bead clean up step should be utilised if g-Tube fragmentation is chosen.

Overall based on our investigation it appears that g-Tubes can provide reasonably consistent DNA fragmentation and the fragment size can be varied in a predictable manner by varying the RCF used and the quantity of DNA. To achieve DNA fragments of approximately 20 Kb, for samples ≤ 1µg, 1000 RCF would be appropriate, whereas for 2 µg 1500 RCF would be best.

Samples of 4 µg would likely need an RCF between 1500 and 2000 as the former resulted in fragments around 25 Kb and the latter around 15 Kb (Table 2).

## Acknowledgements

Our thanks go to Alfred Kedhammar for conducting an effective pilot study which informed our experimental design for this investigation.

---

**Supplementary Figure 1.**
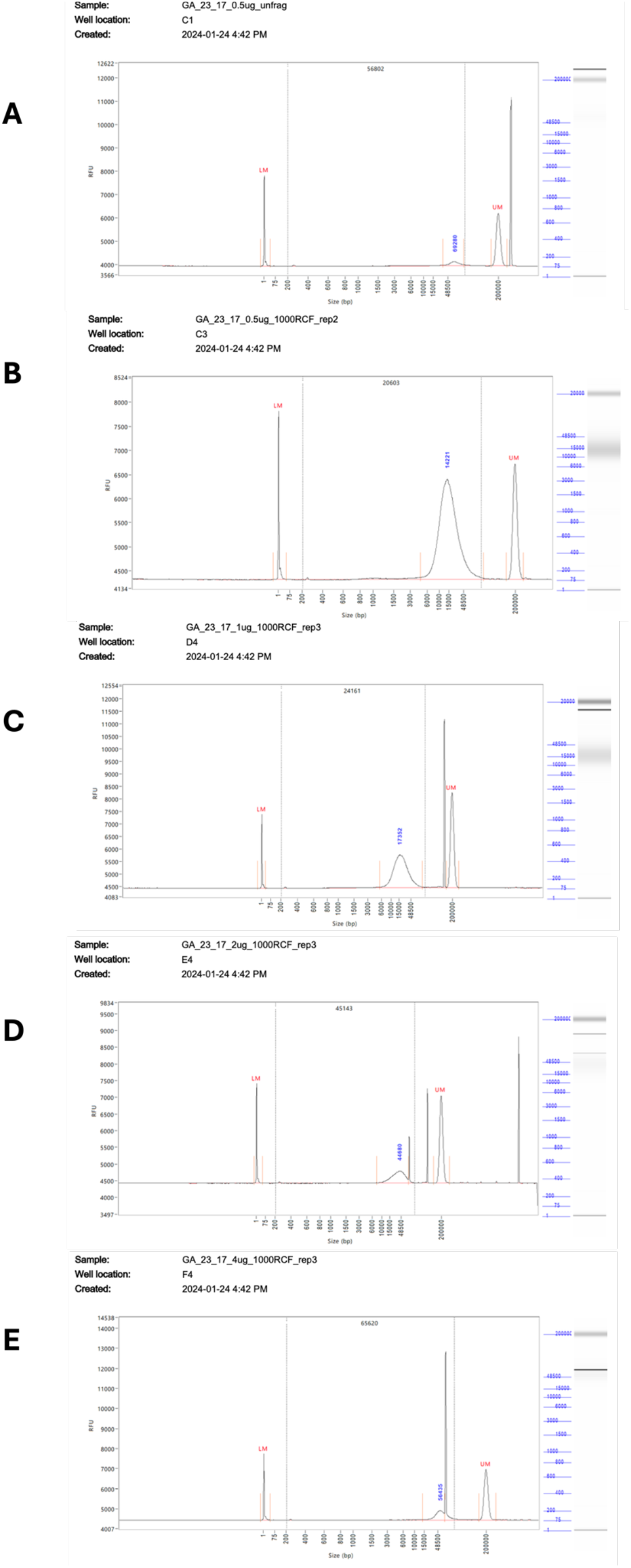
Representative traces of the fragmentation results at RCF 1000 from 0.5 – 4 µg of input DNA. Fragmentation results measured on the 5200 Fragment Analyzer System # M5310AA (Agilent). A) Shows unfragmented DNA, diluted to 0.5 µg / 50 µL. B) Shows DNA diluted to 0.5 µg / 50 µL fragmented at RCF 1000. C) Shows DNA diluted to 1 µg / 50 µL fragmented at RCF 1000. D) Shows DNA diluted to 2 µg / 50 µL fragmented at RCF 1000. E) Shows DNA diluted to 4 µg / 50 µL fragmented at RCF 1000.

